# Pretreatment with LCK Inhibitors Chemosensitizes Cisplatin-Resistant Endometrioid Ovarian Tumors

**DOI:** 10.1101/2020.06.18.154732

**Authors:** Katie K. Crean-Tate, Chad Braley, Goutam Dey, Emily Esakov, Caner Saygin, Alexandria Trestan, Daniel J. Silver, Soumya M. Turaga, Elizabeth Connor, Robert DeBernardo, Chad M. Michener, Peter G. Rose, Justin Lathia, Ofer Reizes

**Affiliations:** Department of Gynecologic Oncology, Cleveland Clinic Foundation, Women’s Health Institute, Cleveland, OH; Department of Cardiovascular and Metabolic Sciences, Lerner Research Institute, and Case Comprehensive Cancer Center, Cleveland, OH; Department of Internal Medicine, The Ohio State University, Columbus, OH

## Abstract

**Objective:** To evaluate LCK inhibitors (LCKi) as chemosensitizing agents for platinum-resistant endometrioid ovarian carcinoma.

**Methods:** KM Plotter survival data was obtained for endometrioid ovarian cancer based on LCK mRNA expression. Cisplatin resistant endometrioid ovarian carcinoma cell lines were cultured and treated first with LCKi or vehicle, then combination LCKi-cisplatin. Cell viability was assessed via CellTiter-Glo, and apoptosis with Caspase 3/7 Assay. Protein lysates were isolated from treated cells, with *γ*-H2Ax, a DNA adduct marker, assessed. *In vivo* study compared mice treated with vehicle or LCK inhibitor followed by LCK inhibitor, cisplatin, or combination therapy. One-way ANOVA and two sample t-test were used to assess statistical significance with GraphPad Prism.

**Results:** KM plotter data indicated LCK expression is associated with significantly worse median progression-free survival (HR 3.19, p=0.02), and a trend toward decreased overall survival in endometrioid ovarian tumors with elevated LCK expression (HR 2.45, p=0.41). *In vitro*, cisplatin resistant ovarian endometrioid cells treated first with LCKi followed by combination LCKi-cisplatin treatment showed decreased cell viability and increased apoptosis. Immunoblot studies revealed inhibition of LCK led to increased expression of *γ*-H2AX. *In vivo* results demonstrate treatment with LCKi followed by LCKi-cisplatin leads to significantly slowed tumor growth.

**Conclusions:** We identified a targetable pathway for chemosensitization of platinum resistant endometrioid ovarian cancer with initial treatment of LCKi followed by co-treatment with LCKi-cisplatin.

## 1. Introduction

Gynecologic malignancy is common, with endometrial and ovarian cancers the most common types in the United States. Ovarian cancer is the most fatal gynecologic malignancy in the United States, with only a 48% survival at 5 years after diagnosis[1]. Typically, advanced disease in ovarian cancer is treated with cytoreductive surgery and platinum-based chemotherapy. Up to 15% of ovarian cancers have endometrioid subtype histologically[2]. Unfortunately, high-grade endometrioid cancers prove difficult to treat due to recurrence and chemoresistance[3]. In ovarian cancer, while up to 85% of patients will enter remission with standard treatment of debulking surgery and platinum-taxane chemotherapy, most of these patients will recur[4]. In the 15% of patients failing standard therapy, disease persists or progresses within the first six months after chemotherapy, indicating platinum-resistant disease. For those who do enter remission, progression to platinum-resistant disease is pervasive[5]. The prognosis is particularly poor in those with platinum-resistant disease, with response rates below 20% for subsequent lines of chemotherapy and continued decrease in disease free interval with each subsequent therapy[6]. As such, recurrent ovarian cancer is known as incurable, with goals of care aimed at symptom management with alternative regimens of chemotherapy[7]. Given the poor prognosis in those with platinum-resistant disease, identification of pathways of chemoresistance and subsequent chemosensitive therapies are on the forefront of cancer treatment[8].

Endometrioid tumors have previously been shown to exhibit a self-renewing population of cells termed cancer stem cells (CSC). CSCs are associated with both tumor recurrence and chemoresistance in multiple tumor types[9]–[12]. CD55, a membrane complement regulatory protein, has previously been indicated as a useful marker for high grade, resistant tumors in many tumor types [13]–[16]. CD55 is highly expressed in endometrioid CSCs [13]. Saygin et al. determined that patient with endometrioid tumors highly expressing CD55, exhibit a poorer progression free survival compared to patients with tumors with low expression, and that CD55 is highly expressed in platinum-resistant ovarian endometrioid cancer cells. Saygin et al. also showed that CD55 is necessary for maintenance of chemoresistance via activation of the non-receptor tyrosine kinase Lymphocyte Cell-Specific Protein-Tyrosine Kinase (LCK) to enhance cisplatin resistance by inducing DNA repair genes, a novel signaling pathway [13]. *In vitro* studies using saracatinib, an investigational drug that inhibits the LCK pathway, show sensitization of cisplatin resistant endometrioid cells to cisplatin in a CSC population. With inhibition of LCK, DNA repair genes were attenuated and cancer cells then showed increased sensitivity to cisplatin. Phase I studies of saracatinib utilized in multiple cancer types have indicated an appropriate safety profile, though follow up randomized human trials in ovarian cancer have fallen short in translating the effect into clinical use[17], [18]. With our initial results and proposed mechanism of action of chemosensitization, we hypothesized that treating cancer cells first with an LCK inhibitor followed by co-treatment with an LCK inhibitor and cisplatin would enhance the chemosensitization effect. Our primary objective was to test this hypothesis *in vitro*, followed by *in vivo* proof of concept test in cisplatin resistance endometrioid cancer model.

## 2. Methods

### 2.1 Cell Culture

Ovarian endometrioid adenocarcinoma cell lines A2780 (cisplatin sensitive) and its cisplatin resistant daughter cell line CP70 were cultured in DMEM medium supplemented with 10% heat-inactivated fetal bovine serum at 37°C in a humidified atmosphere in 5% CO_2_. Cisplatin resistant endometrioid cancer cell line HEC1a was cultured in modified McCoy’s 5a medium supplemented with 10% heat-inactivated fetal bovine serum, also at similar conditions. Cell lines were obtained from Cleveland Clinic centralized research core facility, through which cell lines were previously obtained from the American Type Culture Collection (ATCC) and authenticated. At approximately 80% confluence, trypsin (0.25%)/EDTA solution or Accutase was used to lift cells for passaging as needed for continued experiments until passage 10, at which point a fresh allotment of cells was plated. Cisplatin was obtained from Cleveland Clinic Hospital pharmacy, with 1mg/mL stock solutions stored at room temperature protected from light given its photosensitivity. Saracatinib (AZD0530) was purchased from Selleck Chemicals and 10 *μ*M stock solutions were aliquoted and stored at −20°C. PP2 (AG1879) was purchased from Selleck Chemicals and 10 *μ*M stock solutions aliquoted and stored at −20°C.

### 2.2 Proliferation Assays and Caspase 3/7 Assays

The appropriate cancer cells for each experiment were pre-treated with Saracatinib (1*μ*M), PP2 (10-50 *μ*M), or vehicle (DMSO at similar concentration to drug of interest) for 4 days in T75 flasks. Cells were then plated in 96-well plates at 5,000 cells/well on seeding Day 0, manually counted by hemocytometer using Trypan blue dye exclusion as live cell marker. Cisplatin was then applied the next day at doses of 0-10 *μ*M, with/without Saracatinib, PP2 or vehicle, and treatment was ongoing for 4 to 6 days. Measured proliferation was assessed by CellTiter-Glo (Promega, Southampton, UK) as per manufacturer’s instructions. Percentage survival was normalized to the untreated control for each group.

Caspase 3/7 Assay kit (Promega, Southampton, UK) was utilized to assess apoptosis as per manufacturer’s instructions. This was performed alongside CellTiter-Glo to correct for viable cell density. Relative caspase activities were normalized to untreated controls in each group, with activity assessed from 30 - 120minutes.

### 2.3 Immunoblotting

Protein lysates were obtained with cell lysis in 20mM Tris-HCl (pH 7.5), 150mM NaCl, 1 mM Na2EDTA, 1% NP-40, 1 mM EGTA, 1% sodium pyrophosphate, 1 mM β-glycerophosphate, 1mM sodium orthovanadate, 1 *μ*g/mL leupeptin, 20 mM NaF and 1 mM PMSF. Protein concentrations were measured with BCA Protein Assay Kit (ThermoFisher Scientific). Protein concentrations from 20-50 *μ*g of total protein were resolved in 10-12% SDS-PAGE and transferred to PVDF membrane. Membranes were incubated overnight at 4°C with primary antibodies against pLCK (Y394) (1:1000) (R&D Systems), GAPDH (1:1000) (Cell Signaling), and *γ*-H2AX (1:1000) (Cell Signaling). Secondary anti-mouse or anti-rabbit IgG antibodies conjugated to horse radish peroxidase (HRP) (1:3000) (Cell Signaling) or (1:25,000) (ProMega) were used. ECl was then used (Pierce) to visualize immunoreactive bands.

### 2.4 In vivo Study

All animal procedures were evaluated and approved prior to initiation by the Institutional Animal Care and Use Committee (IACUC) of the Cleveland Clinic Lerner Research Institute. NOD severe combined immunodeficient (SCID) IL2R gamma (NSG) mice were purchased from the Biological Response Unit (BRU) at the Cleveland Clinic and housed in microisolator units under IACUC protocol #2018-1940. Thirty mice were injected intraperitoneally with 1 million CP70-luciferase virally transduced cells. At the time of injection (day 0), mice were placed in one of two arms, which started day 3: six mice began receiving pre-treatment with Saracatinib (Selleck), 25mg/kg dissolved in 0.5% hydroxypropyl methylcellulose (Sigma-Aldrich), 0.1% Tween 80 (Sigma-Aldrich) via oral gavage three days per week, and 24 mice received vehicle via oral gavage on the same schedule.

Bioluminescence images to detect tumor burden were taken with Xenogen *in vivo* imaging system (IVIS, PerkinElmer) using D-luciferin as previously described[19]. Mice received an IP injection of D-luciferin (Goldbio LUCK-1G, 150mg/kg in 150*μ*L) under inhaled isoflurane anesthesia. Images were analyzed (Living Image Software) and bioluminescence plots of photon flux (photons/second/cm^2^/steradian) over time were computed for each mouse, with normalization against day 0 signal values. Non-tumor and black backgrounds were also subtracted from each tumor burden region of interest. All images were obtained with a 15 second exposure. On day 16 when all mice were confirmed to have tumor by IVIS, mice pretreated with saracatinib were also treated with cisplatin (2.5mg/kg, 3 times per week) injected intraperitoneally. On day 16, mice pretreated with vehicle were randomly assigned to one of four arms (6 mice per arm), whereby they were treated with cisplatin, saracatinib, combination cisplatin and saracatinib, or vehicle alone. Mice were sacrificed on day 30 and all visible tumor was collected for future studies. All mouse procedures were performed under adherence to protocols approved by the Institute Animal Care and Use Committee at the Lerner Research Institute, Cleveland Clinic.

### 2.5 Statistical Analysis

Statistical analysis was calculated by one-way ANOVA and two sample t-test, with p-values included. Statistical significance is denoted via * to represent p-value of <0.05 but >0.01, ** representing p-value of <0.01 but >0.001, and ** representing p-value <0.001. For proliferation assays, IC50 was calculated using nonparametric values set to nonlinear fit curve as per statistical analysis performed with GraphPad Prism. Survival data was obtained from Kaplan-Meier Plotter (KM Plotter: http://kmplot.com/analysis/) for endometrioid ovarian cancer based on CD55 and LCK mRNA expression. KM Plotter survival data is obtained from an online database collected from The Cancer Genome Atlas (TCGA), Gene Expression Omnibus (GEO) and European Genome-phenome Archive (EGA).

## 3. Results

### 3.1 CD55 and LCK expression are associated with poorer patient survival

Given the previously described mechanism of CD55 regulation of cisplatin resistance via the LCK pathway, we hypothesized that both increased CD55 and LCK expression would be associated with worse clinical outcomes. We assessed survival outcomes with increased CD55 and LCK expression for endometrioid ovarian cancer using Kaplan-Meier Plotter database (KM Plotter: http://kmplot.com/analysis/). CD55 expression is associated with a significantly worse median progression-free survival (HR 2.98, p<0.05, **Fig. 1a**) but did not inform median overall survival (HR 6.48, p=0.055, **Fig. 1b**). We then assessed survival outcome in LCK expressed ovarian tumors. In endometrioid ovarian cancer, LCK expression is associated with significantly worse median progression-free survival (HR 3.19, p=0.02, **Fig. 1c**). Overall survival is not significantly different between groups, though a non-significant trend toward decreased survival was seen with HR 2.45 (p=0.41, **Fig. 1d**). These data indicate increased CD55 and LCK expression in endometrioid ovarian cancer correlates with poorer clinical outcomes.

**Figure 1.**
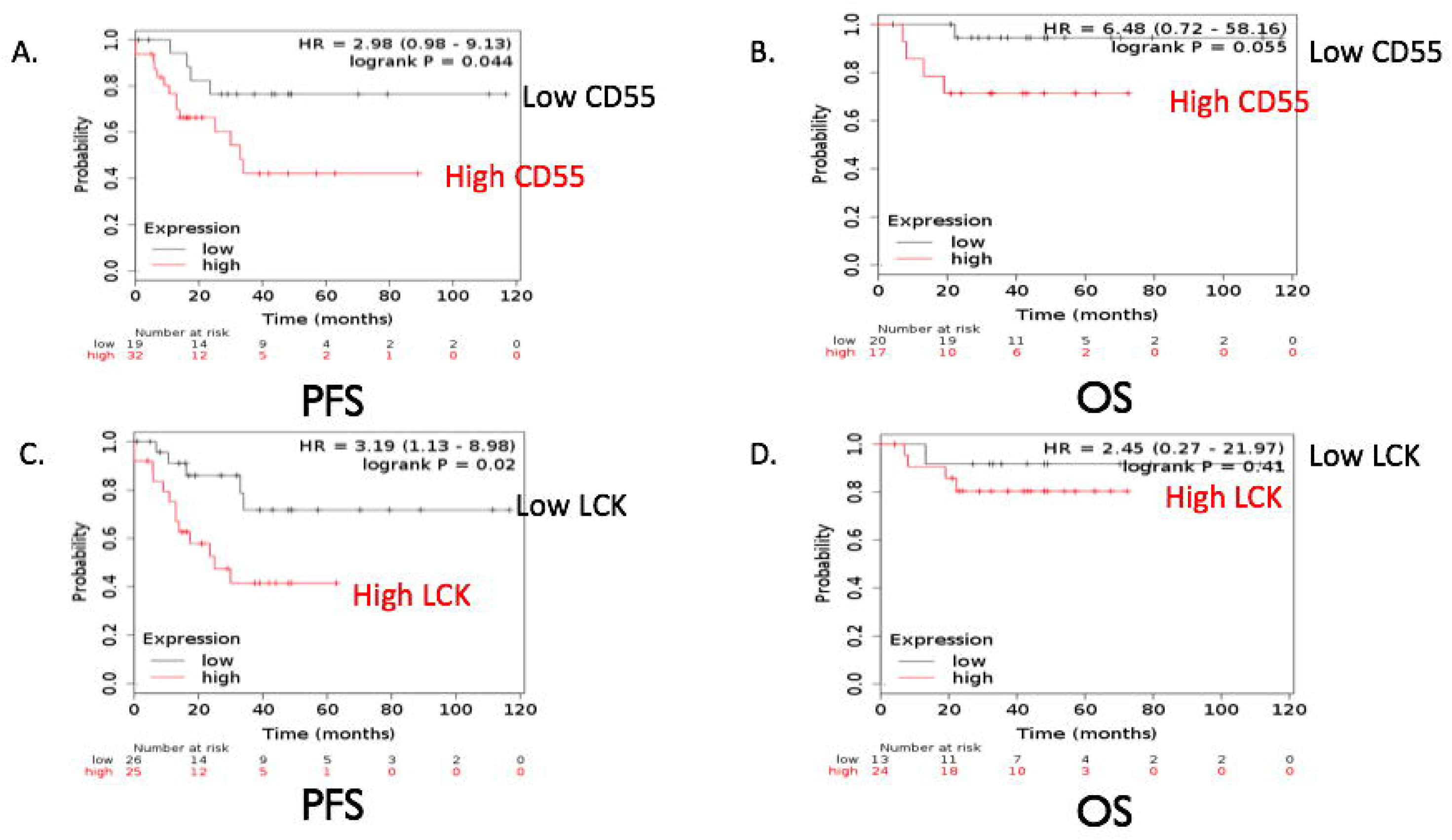
CD55 and LCK expression are associated with poorer patient survival. Kaplan-Meier progression-free and overall survival curves were obtained from Kaplan-Meier Plotter (KM Plotter: http://kmplot.com/analysis/) for endometrioid ovarian cancer patients who had high versus low tumor mRNA expression of CD55 **(A, B)** and LCK **(C, D)** prior to therapy.

### 3.2 Pretreatment with LCK inhibitors chemosensitize cisplatin resistant endometrioid cells and increase apoptosis

In our prior studies, LCK inhibition in the ovarian endometrioid CSC population led to increased cisplatin sensitivity[13]. Given that CSCs are known to be a chemoresistant population closely associated with disease recurrence, we theorized that inhibition of the LCK pathway would lead to sensitization of platinum resistant endometrioid cells. To test our hypothesis, we performed *in vitro* cellular proliferation assays in cisplatin resistant ovarian endometrioid cells (CP70) treated with vehicle or the LCK inhibitor (LCKi) saracatinib followed by cisplatin sensitivity with varying concentrations of cisplatin. It is noted that these tests would not include only a CSC population as with Saygin’s study[13], but rather included all CP70 cells. As there was no significant difference with co-treatment of LCKi and cisplatin alone, we hypothesized that pretreatment with LCKi is required to effectively sensitize these cells to cisplatin. We found that CP70 cells demonstrated significantly decreased proliferation when pretreated with saracatinib and then treated with combination saracatinib-cisplatin (**Fig. 2a**). We also tested these cells in a Caspase Glo assay to assess for apoptosis. We determined that CP70 cells pretreated with saracatinib followed by cisplatin plus saracatinib increased apoptosis compared to no pretreatment (**Fig. 2b**). These findings were replicated in HEC1a, a cisplatin resistant endometrioid cancer model. HEC1a pretreated with saracatinib followed by cisplatin cotreatment with saracatinib exhibited inhibition of cell viability (**Fig. 2c**) and increased apoptosis (**Fig. 2d**) compared to vehicle treated cells.

**Figure 2.**
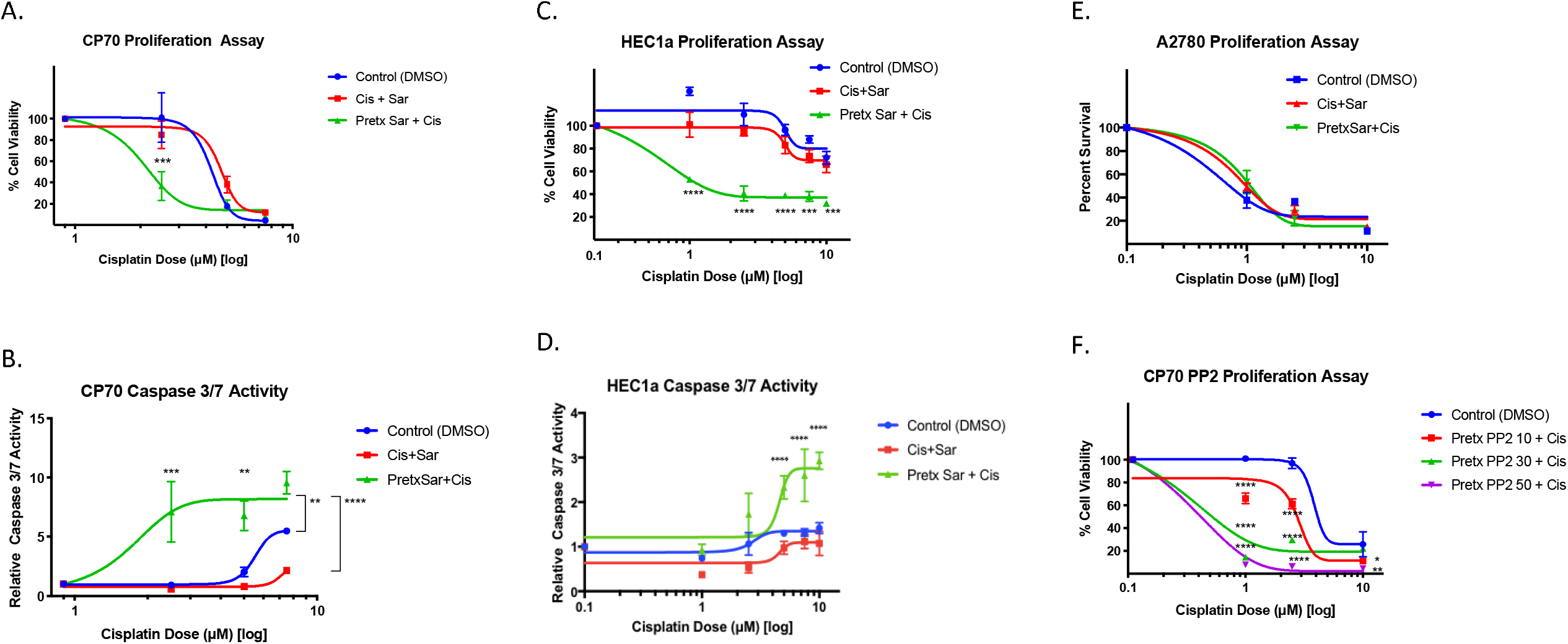
LCK inhibitors chemosensitize cisplatin resistant endometrioid cells and increase apoptosis. Cisplatin resistant ovarian endometrioid cells (CP70) were cultured and pretreated with an LCK inhibitor (saracatinib) or vehicle, followed by vehicle or combination LCKi-cisplatin, followed by cell viability assay performed with the CellTiterGlo Assay **(A)**. Caspase 3/7 Assay was then performed to assess apoptosis **(B).** A second cisplatin resistant endometrioid cell line (HEC1a) was similarly treated and tested with subsequent proliferation and apoptosis assays performed **(C, D).** Cisplatin sensitive ovarian endometrioid cells (A2780) were cultured and treated according to the aforementioned paradigm **(E).** An alternative LCK inhibitor (PP2) was utilized for pretreatment in CP70 cells followed by co-treatment with PP2-cisplatin **(F).** All data represent at minimum three independent experiments.

To validate our hypothesis that initial LCK inhibitor treatment followed by co-treatment of LCKi-cisplatin leads to decreased proliferation specifically in cisplatin resistant endometrioid cells, we performed a proliferation assay in A2780 cells, the chemo-naive parent cell line of CP70. We found that pretreatment with saracatinib, as well as simultaneous treatment with saracatinib and cisplatin, did not significantly alter the cell viability (**Fig. 2e**). To validate our hypothesis that saracatinib chemosensitizes these cells via LCK inhibition, we pretreated CP70 cells with an alternative LCK inhibitor, PP2, followed by cisplatin co-treatment, and found a dose responsive decrease in cell proliferation seen as compared to control groups treated with cisplatin only (**Fig. 2f**). These data indicate that platinum resistant endometrioid cells pretreated with LCKi followed by co-treatment with LCKi and cisplatin show decreased cell proliferation and increased apoptosis.

### 3.3 Cisplatin resistant endometrioid cells treated with LCK inhibitors reveal DNA double strand breaks

To investigate the mechanism by which LCK inhibitors decrease cisplatin resistance, we tested whether LCK inhibitors decrease phosphorylation of LCK in CP70 cells by immunoblot. We found that phosphorylated LCK (pLCK) was significantly reduced when cells are exposed to LCK inhibitors saracatinib and PP2. GAPDH was used as a loading control (**Fig. 3a**). We tested for DNA double strand breaks by blotting for *γ*H2AX, a histone that is phosphorylated after DNA double strand breaks occur [20], [21]. We found an increase in *γ*H2AX in LCK-inhibitor treated cells (**Fig.3b**). These data indicate that endometrioid ovarian cancer cells exposed to LCKi demonstrate a decrease in phosphorylated LCK and an increase in DNA double strand breaks.

**Figure 3.**
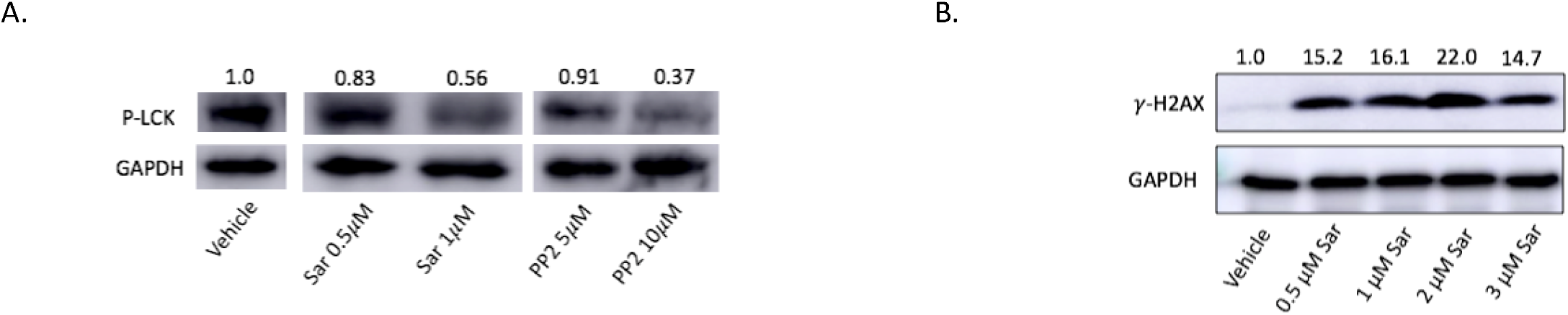
Cisplatin resistant endometrioid cells treated with LCK inhibitors indicate decreased pLCK and ovarian endometrioid cells treated with LCK inhibitor indicate increased DNA double strand breaks. Protein lysates from cisplatin resistant ovarian endometrioid cancer cells (CP70) treated with vehicle (DMSO), LCK inhibitor saracatinib (Sar), or PP2 were immunoblotted for pLCK, with GAPDH used as a loading control. Values normalized to vehicle control. **(A)** Protein lysates from ovarian endometrioid cancer cells (TOV112D) treated with varied doses of saracatinib (Sar) were immunoblotted for *γ*H2AX, with GAPDH used as a loading control **(B)**.

### 3.4 Treatment with LCK inhibitor followed by co-treatment of LCKi-cisplatin decreases tumor growth in vivo

Given the *in vitro* findings, we next tested our hypothesis in an *in vivo* model. We injected NSG mice with CP70 cells virally transduced with luciferase and utilized *in vivo* imaging system (IVIS) to assess weekly tumor growth. After injection of tumor cells (day 0), mice were placed in one of two arms: pre-treatment with saracatinib or vehicle via oral gavage three days per week, initiated day 3. On day 16 when all mice were confirmed to have detectable tumor on IVIS, mice pretreated with saracatinib were injected with cisplatin three times weekly, and mice pretreated with vehicle were randomly assigned to one of four arms: cisplatin, saracatinib, combination cisplatin and saracatinib, or vehicle alone, given three times per week. Mice were then sacrificed on day 30 (**Fig. 4a**). Mice treated first with vehicle followed by cisplatin, saracatinib, or combination cisplatin and saracatinib all showed a steady increase in tumor burden over time. However, in mice first treated with saracatinib followed by combination cisplatin and saracatinib therapy, tumor growth appeared stable or attenuated (**Fig. 4b, c**). At the experimental endpoint (Day 30), the vehicle group indicated that tumor growth was statistically similar to cisplatin, saracatinib, and combination cisplatin and saracatinib arms. However, there was a significant reduction in tumor growth seen in the pretreatment saracatinib followed by combination saracatinib-cisplatin arm as compared to vehicle as well as combination only (**Fig. 4d**). These data demonstrate that pretreatment with LCKi followed by LCKi-cisplatin co-treatment leads to decreased tumor burden in cisplatin resistant endometrioid ovarian cancer *in vitro*.

**Figure 4.**
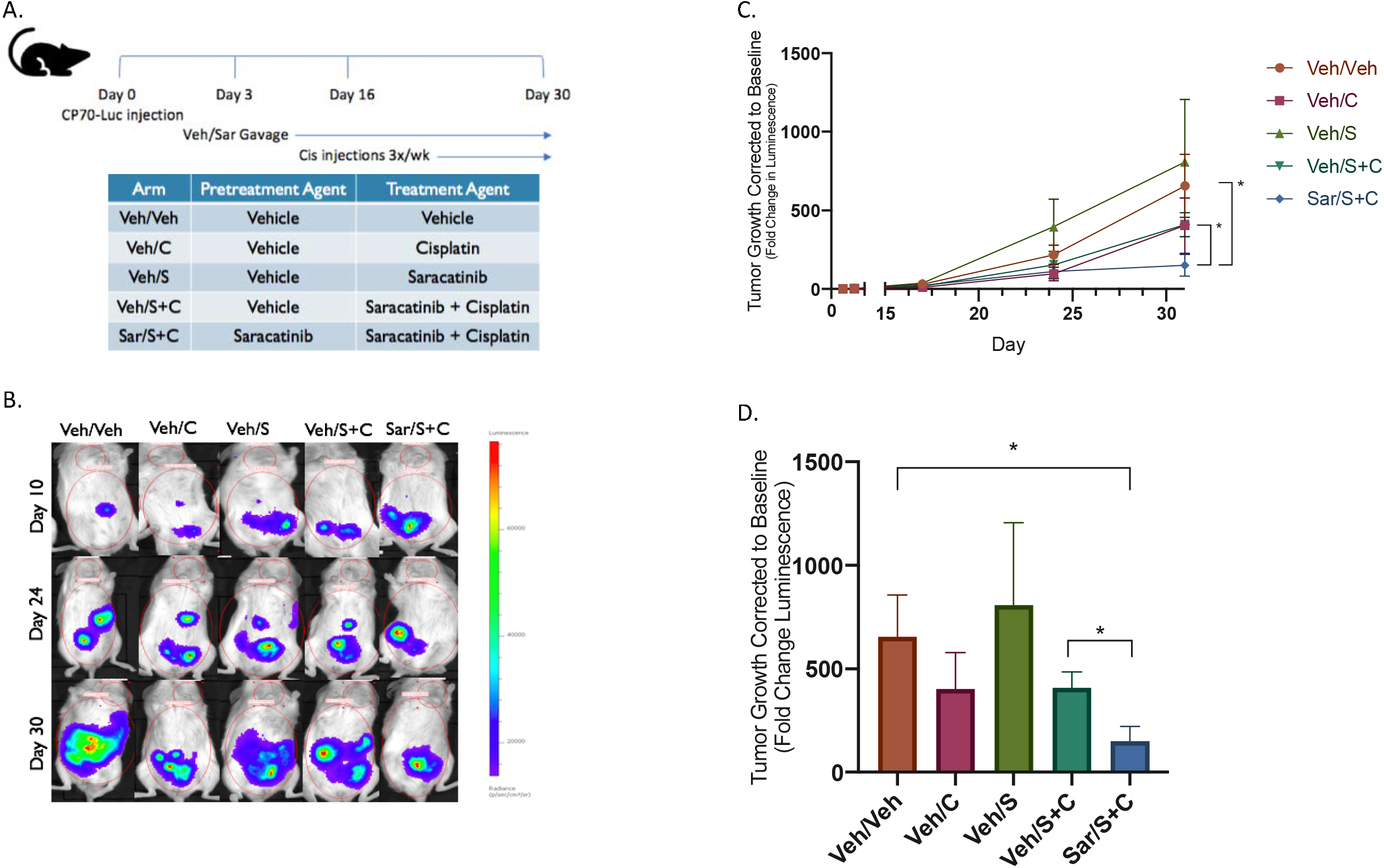
Pretreatment with LCK inhibitor followed by LCKi-cisplatin treatment attenuates tumor burden. NSG mice were injected with CP70-luciferase transfected cells followed by pretreatment with LCKi (6 mice) or vehicle (24 mice) for 14 days. LCKi mice were then co-treated with LCKi and cisplatin, and vehicle mice were randomized to further treatment with vehicle, cisplatin, saracatinib, or combination (6 mice per arm) **(A)**. IVIS imaging was obtained on a weekly basis to assess tumor growth **(B)**. IVIS luminescence was corrected to baseline for each arm and assessed over time **(C)** and at the experimental endpoint **(D)**.

### 3.5 LCK inhibition as a targetable pathway for platinum resistant ovarian endometrioid cells

It has been established that downstream of CD55, LCK stimulates expression of DNA repair genes, leading to cisplatin resistance[13]. We hypothesized that inhibition of LCK would lead to sensitization of cisplatin resistant cells. We found that pretreating cisplatin resistant ovarian endometrioid cells with an LCK inhibitor prior to co-treatment of LCKi and platinum therapy leads to sensitization of a chemoresistant tumor. This identifies a targetable pathway and indicates LCK inhibitors may play a role in adjunctive therapy for platinum resistant ovarian endometrioid cancer **(Fig. 5).**

**Figure 5.**
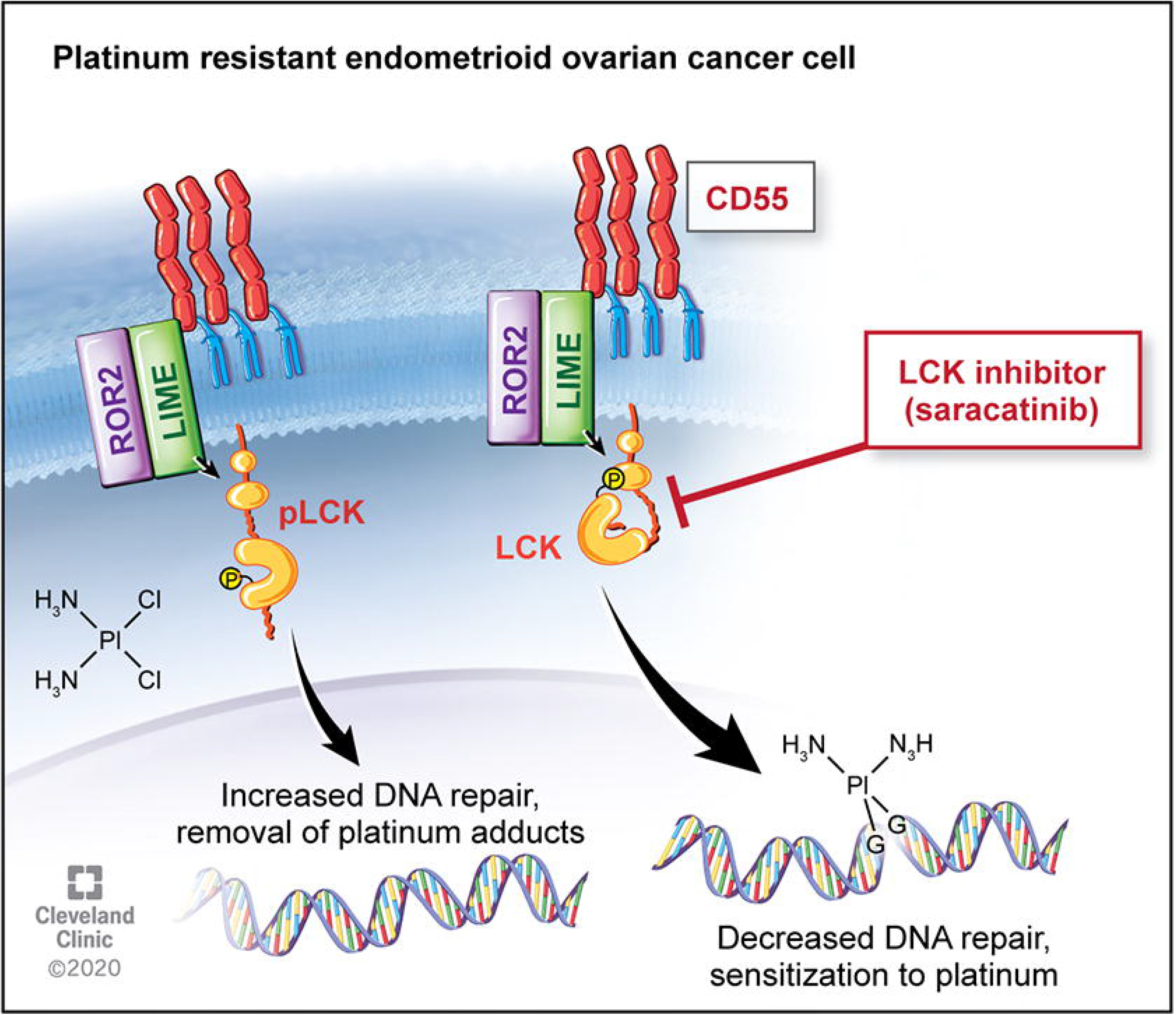
LCK pathway regulates cisplatin resistance in endometrioid tumors. Downstream of CD55, LCK stimulates expression of DNA repair genes, leading to cisplatin resistance. This targetable pathway identifies LCK inhibitors as adjunctive therapy for platinum resistant ovarian endometrioid cancer.

## 4. Discussion

Despite most ovarian cancers displaying an excellent initial response to standard chemotherapy, the majority of advanced stage patients recur, with eventual resistance to our most effective chemotherapy agents. This pattern of pervasive relapse and ensuing chemoresistance is the cause for the poor survival rates seen in ovarian cancer today [3], [5], [9]. Studies have focused on cell populations known for chemoresistance, cancer stem cells (CSCs), in order to identify a targetable pathway to reduce recurrence[9]–[11]. Previous studies by Saygin et al.[13] identified a novel pathway leading to chemoresistance in endometrioid tumors in which CD55 mediated DNA repair via phosphorylation of LCK. We assessed clinical outcomes associated with LCK expression. We found that endometrioid ovarian cancers highly expressing CD55 indicated worse clinical outcomes, with significant effects in progression free survival (PFS) (HR 2.98, p=0.04). Given our prior finding that the CD55/LCK pathway is involved in chemoresistance, we assessed survival in tumors highly expressing LCK. We found that high LCK expression predicted an even more significant effect on PFS with a three-fold increase in survival from 13 months to 34 months in low vs high LCK expressing tumors. This data suggest a clinical benefit to addressing tumors with increased LCK expression, and thus a potential targetable pathway in recurrent ovarian endometrioid tumors.

Standard chemotherapy in ovarian cancer includes a platinum and taxane agent, and survival decreases as response to platinum therapy diminishes. Prior studies have found that cisplatin resistance is seen with multiple pathways, including increased DNA repair enzyme expression and associated reduction in DNA adducts [9], [13], [22]. Through LCK inhibition, DNA repair enzyme expression is attenuated. Given this anticipated initial chemosensitization step, we pursued pretreatment with LCK inhibitors followed by co-treatment with cisplatin, and found that this technique was effective in decreasing cancer cell populations and increasing apoptosis in vitro (**Fig. 2**). We verified the effects of inhibition of LCK on DNA damage, and found an increase in DNA adduct formation with LCK inhibition in immunoblot studies (**Fig. 3**), an indication that targeting this pathway allows platinum therapy to function in a previously cisplatin resistant cell population.

A common challenge in translational research is that while *in vitro* studies may prove promising, translating this to effective *in vivo* studies and clinical trials can prove difficult. Saracatinib, an investigational LCK inhibitor, has been studied for many cancer types, with mixed results. Studies on safety found appropriate dosing for saracatinib in humans for effective pharmacodynamics while limiting toxicity, indicating this drug would be tolerable in clinical trials [17]. While utilizing saracatinib as monotherapy has not proven efficacious, combination therapy has yielded more promising results. In a study combining saracatinib with carboplatin and/or paclitaxel in solid tumors, objective responses were seen in ovarian, breast and skin cancers, with longest response durations seen in patients with ovarian cancer [17]. However, a randomized trial further assessed treatment with saracatinib in combination with weekly paclitaxel in platinum-resistant ovarian cancer, and found that co-treatment of saracatinib with weekly paclitaxel did not improve outcomes [18]. Of note, the majority of these tumors were serous histology, and patients received only weekly Taxol without platinum in addition to saracatinib. There is no clinical randomized data assessing saracatinib with cisplatin use in platinum resistant patients. We see from this clinical data that saracatinib is well-tolerated and may have a role in combination therapy in platinum resistant disease. We tested this hypothesis with a novel administration of saracatinib followed by co-treatment with cisplatin and found decreased rate of tumor growth *in vivo* (**Fig. 4**), identifying a targetable pathway **(Fig. 5)** and providing a novel therapeutic regimen for platinum resistant ovarian endometrioid carcinoma.

This study’s strength lies in the proof of concept findings using both *in vitro* and *in vivo* models. Additionally, whereas prior studies focused on a specific cell population, cancer stem cells, this study utilized a more heterogenous cell population, more closely simulating a typical tumor microenvironment. Further investigation should be performed in additional histologic types such as serous and clear cell, as well as substitution of cisplatin for co-treatment with carboplatin, a commonly used platinum agent. Patients with platinum resistant disease have often received multiple lines of chemotherapy previously and further treatments offer limited therapeutic benefit. Given our promising findings, further studies are indicated to pursue LCK inhibitors as an adjunctive therapy to platinum resistant disease in clinical trials.

## Conclusion

In summary, we identified a targetable pathway for chemosensitization of platinum resistant ovarian endometrioid cancer. We found that pretreatment with LCK inhibitors followed by cotreatment with cisplatin leads to decreased cell viability and increased apoptosis *in vitro*. This is associated with increased DNA adduct formation and significantly reduced tumor growth *in vivo*. Further studies are needed to assess the mechanisms behind the enhanced efficacy of pretreatment, as well as further investigation of LCK inhibitors as adjunctive therapy for platinum resistant endometrioid ovarian carcinoma, including other histological subtypes.

## Abbreviations

LCK: Lymphocyte Cell-Specific Protein-Tyrosine Kinase

## Acknowledgements

Research in the Reizes laboratory is supported by VeloSano Bike to Cure Impact Award, the Laura J. Fogarty Endowed Chair for Uterine Cancer Research and the Cleveland Clinic Foundation Lerner Institute.

## Author Contribution

KKCT: Hypothesis generation, study design, experiment execution, data analysis, and primary manuscript authorship.

CB, GD: Contributed to experiment execution and data collection.

EE: Contributed to data organization, analysis, and manuscript editing.

CS, EC: Contributed to hypothesis generation, experiment execution and data organization.

AT: Contributed to experiment execution and data collection.

DS: Contributed to data organization and manuscript editing.

ST: Contributed to experiment execution.

RB, CM, PGR: Contributed to data and manuscript review.

JL: Contributed to hypothesis generation and study design.

OR: Significant contributions to hypothesis generation, study design, and manuscript editing.

